# MACGA: Multi-scale Adaptive Convolution with Graph Attention for LncRNA–Disease Association Prediction

**DOI:** 10.1101/2025.07.14.664702

**Authors:** Xiulai Li, Xin Li, Wen Zhang, Wei Wu

## Abstract

Accurate prediction of lncRNA-disease associations (LDAs) is crucial for understanding complex disease mechanisms and advancing precision medicine. Existing computational methods based on graph neural networks (GNNs) often suffer from over-smoothing issues, limiting their predictive accuracy. To address this challenge, we propose MACGA, a novel neural network framework integrating Multi-scale Adaptive Convolution with Graph Attention networks. MACGA effectively captures both local neighborhood information and global high-order structural features by combining PCA-based dimensionality reduction, local node feature extraction via graph attention mechanisms, and multi-scale feature aggregation through adaptive convolution. Extensive experiments using five-fold cross-validation demonstrated that MACGA significantly outperformed four representative state-of-the-art methods (AUC=0.9463, AUPR=0.4119). Case studies on prostate and colon cancers further validated the reliability of MACGA, with top-ranked lncRNA predictions well-supported by existing experimental literature. Moreover, we extended the proposed framework to predict disease-drug associations via lncRNA-bridging strategy, highlighting its potential for drug repositioning. Overall, MACGA provides a powerful computational tool for accurately predicting lncRNA-disease associations and exploring novel therapeutic opportunities. The source code and dataset used in this paper are available at: https://github.com/huifeideyu518/MACGA.

## 1. Introduction

Non-coding RNAs (ncRNAs) represent a diverse class of RNA molecules that, despite lacking protein-coding capacity, play crucial regulatory roles in gene expression across a range of biological processes, including chromatin remodeling, transcription, RNA splicing, and translation [1]. Among the various ncRNA subtypes, long non-coding RNAs (lncRNAs)-defined as transcripts exceeding 200 nucleotides in length-have garnered substantial attention due to their key involvement in numerous complex diseases, including cancers, cardiovascular disorders, and neurological conditions [2,3]. Recent investigations have elucidated the functional significance of specific lncRNAs in disease pathogenesis. For instance, lncRNA CCAT1 has been shown to promote prostate cancer cell proliferation, migration, and invasion through modulation of the miR-490-3p/FRAT1 signaling axis [4]. Similarly, lncRNA LINC00942 contributes to chemoresistance in gastric cancer cells by inhibiting MSI2 degradation, thereby stabilizing c-Myc mRNA [5]. These findings underscore that identifying potential lncRNA-disease associations not only enhances our mechanistic understanding of diseases but also reveals novel biomarkers and therapeutic targets for clinical application. However, the systematic experimental identification of lncRNA-disease associations remains labor-intensive, cost-prohibitive, and impractical for large-scale screening, highlighting the need for computational approaches to efficiently predict these associations [6].

Current computational methods for predicting lncRNA-disease associations can be broadly categorized into three main approaches: network topology-based methods, matrix factorization and graph autoencoder methods, and deep learning-based graph neural network (GNN) methods [7]. Network topology-based prediction methods typically construct heterogeneous networks and use algorithms such as network diffusion or random walks to infer potential lncRNA-disease associations. For example, Sumathipala et al. [8] proposed a method called LION, which utilizes multi-layer network topology and network diffusion algorithms to predict lncRNA-disease associations for cardiovascular diseases, cancer, and neurological disorders, achieving better performance than similar methods like TPGLDA. Hu et al. [9] proposed the BiWalkLDA algorithm, which constructs an lncRNA-disease network by integrating interaction profiles and gene ontology information via bidirectional random walks (Bi-Random Walks), effectively addressing the cold start problem. Xie et al. [10] introduced the SSMF-BLNP method, which combines selective similarity matrix fusion (SSMF) and bidirectional linear neighborhood label propagation (BLNP) to predict lncRNA-disease associations. Matrix factorization and graph autoencoder methods, on the other hand, optimize prediction results through low-rank decomposition or embedding strategies that integrate multiple similarity matrices. Wu et al. [11] proposed the GAERF method, a classification approach based on graph autoencoders (GAE) and random forests (RF), which effectively identifies lncRNA-disease associations using low-dimensional feature vectors and ensemble learning. Fu et al. [12] developed the MFLDA model, which applies matrix tri-factorization to explore the intrinsic structure of data sources and integrates them using weighted fusion, thereby improving prediction accuracy. In recent years, deep learning-based GNN methods have emerged as powerful tools due to their ability to capture nonlinear relationships and complex network structures. Chen et al. [13] proposed a model called SGGCL, which combines graph neural networks and contrastive learning by generating high-quality positive and negative sample pairs, enhancing prediction accuracy through the use of graph convolutional networks and the RWR algorithm. Shi et al. [14] introduced the end-to-end VGAELDA model, which integrates variational inference and graph autoencoders by alternating between training variational graph autoencoders (VGAE) and graph autoencoders, significantly boosting prediction robustness and accuracy. Li et al. [15] proposed the novel SVDNVLDA model, which combines singular value decomposition (SVD) and node2vec to capture both linear and nonlinear features of lncRNAs and diseases, using the XGBoost classifier to predict potential lncRNA-disease associations. Additionally, Fan et al. [16] proposed the IDHI-MIRW method, which integrates multiple heterogeneous information sources, pointwise mutual information, and the random walk with restart algorithm to construct large-scale lncRNA-disease heterogeneous networks, effectively identifying potential associations. Yao et al. developed the GCNFORMER model, which fuses graph convolutional networks with Transformer architectures, demonstrating superior performance in case studies involving breast cancer, colon cancer, and lung cancer. Although these methods have achieved significant progress in capturing local network topologies, reducing data redundancy and noise, and enhancing model robustness, challenges remain. These include how to comprehensively capture long-range higher-order relationships, how to avoid the inherent over-smoothing issue in deep GNN architectures, and how to better model disease subtype-specific variations. Therefore, future research should focus on further enhancing the overall performance of these methods by strengthening the integration of structural and attribute information and improving both prediction accuracy and robustness while addressing these challenges.

To address these fundamental limitations, we propose an end-to-end hybrid neural network framework specifically designed to capture both local and multi-scale higher-order topological features while simultaneously reducing data redundancy and noise through effective integration of heterogeneous data. Our approach begins with Principal Component Analysis (PCA)-based dimensionality reduction, which eliminates redundant information from high-dimensional lncRNA-disease feature matrices. Subsequently, these denoised features are processed through a Graph Attention Network (GAT) to enable dynamic, adaptive aggregation of local neighborhood information. To enhance the model’s capacity for capturing long-range, higher-order relationships, we incorporate the Multi-scale Adaptive Convolution Module (MACM) from the Multi-scale Graph Clustering Network (MGCN), which adaptively integrates information from 1-hop to k-hop neighbors. Finally, a Multi-Layer Perceptron (MLP) predicts lncRNA-disease association likelihood based on these integrated node embeddings. Through this comprehensive four-step pipeline-denoising, local attention encoding, multi-scale convolutional fusion, and association prediction-our method effectively integrates both structural and attribute information while mitigating the over-smoothing problem inherent in deep GNN architectures, resulting in more discriminative and informative node representations.

The principal contributions of this work are threefold:

**(1) Hybrid neural network architecture** We develop a novel hybrid framework that integrates graph attention mechanisms with multi-scale adaptive convolutions, embedding the MACM module within the GAT architecture to simultaneously capture local and long-range higher-order network structures, thereby effectively alleviating over-smoothing issue prevalent in deeper GNNs.
**(2) End-to-end feature extraction pipeline** We construct a comprehensive pipeline that combines PCA-based denoising, GAT-based local encoding, and MACM-driven multi-scale convolutional fusion, significantly enhancing model robustness and predictive accuracy, particularly for high-dimensional, sparse, and noisy biomedical datasets.
**(3) Cross-scenario application framework** We propose an extensible predictive framework that generalizes from lncRNA-disease associations to lncRNA-drug and disease-drug associations. Using gastrointestinal disease subtypes and related therapeutics as illustrative examples, we demonstrate the broad applicability and translational potential of our method for multi-entity association prediction, offering novel insights and practical strategies for drug repositioning.

The remainder of this manuscript is structured as follows: Section 2 details the materials and methods, including dataset descriptions, preprocessing techniques, and the architectural and training strategies of the proposed neural network framework. Section 3 presents experimental results and analyses, covering evaluation criteria, parameter optimization, performance comparisons, and case studies involving lncRNA-disease associations, disease subtype analysis, and lncRNA-drug/disease-drug predictions. Section 4 concludes with a summary of key contributions, a discussion of limitations, and future research directions.

## 2. Materials and Methods

### 2.1 Method Overview

This paper proposes a hybrid neural network framework that combines Graph Attention Networks (GAT) with the Multi-scale Adaptive Convolution Module (MACM) from Multi-scale Graph Clustering Networks (MGCN) to effectively predict lncRNA-disease associations. The overall framework adopts an end-to-end deep learning architecture, as shown in Figure 1, comprising four main stages: feature dimensionality reduction and denoising, local attention encoding, multi-scale convolution fusion, and association prediction. First, we collect multi-source heterogeneous biological data and construct initial feature representations for lncRNAs and diseases. Subsequently, Principal Component Analysis (PCA) is employed to perform dimensionality reduction on high-dimensional features to eliminate data redundancy and reduce noise interference. Next, we construct a lncRNA-disease bipartite graph and use graph attention networks to dynamically learn local neighborhood representations of nodes. To capture long-range high-order structural relationships, we introduce the multi-scale adaptive convolution module from MGCN, which obtains multi-scale node embeddings by simultaneously aggregating information from 1-hop to *k*-hop neighbors. Finally, the local features encoded by GAT are fused with the multi-scale features extracted by MACM and fed into a multi-layer perceptron for final association probability prediction.

**Fig 1.**
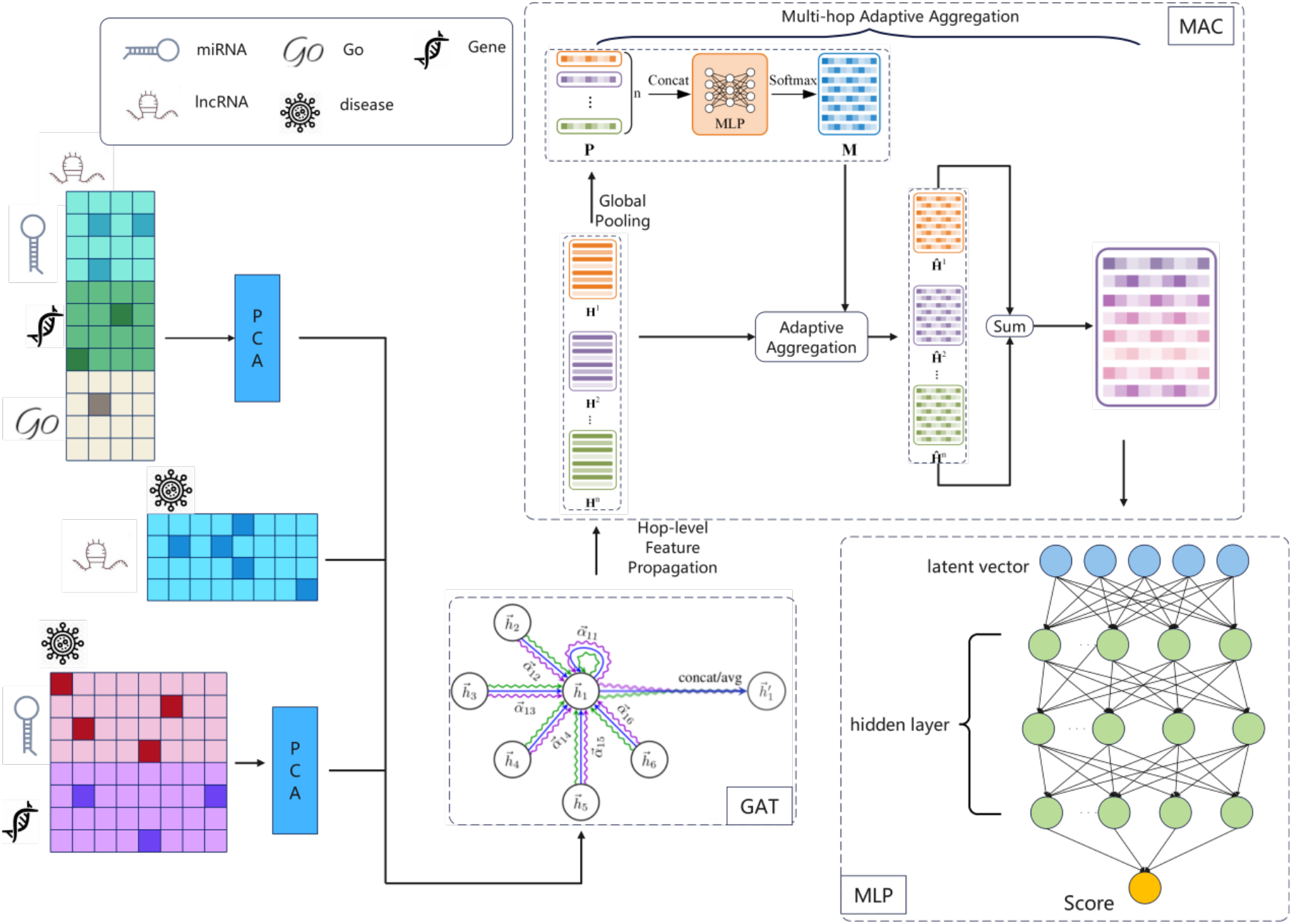
The overall structure of our MACGA for predicting LDA.

### 2.2 Data Collection and Preprocessing

To facilitate the validation of prediction results and avoid using databases with a short time span that may make the results difficult to verify, we collected experimentally validated lncRNA-disease association data from several authoritative biomedical databases that have been published for several years. The primary lncRNA-disease association data were sourced from the LncRNADisease [18], Lnc2Cancer [19], and GeneRIF [20]. After data cleaning and deduplication, we obtained 2,697 lncRNA-disease associations involving 240 lncRNAs and 412 diseases. For lncRNA feature construction, we integrated multi-dimensional biological association information: lncRNA-gene association data were obtained from the LncRNA2Target [21]; lncRNA-GO functional annotation associations were acquired through the GeneRIF [20] and preprocessed using Open Biomedical Annotator; lncRNA-miRNA interaction data were downloaded from the StarBase v2.0 [22]. After data processing, we retained association information for 240 lncRNAs, 5,347 genes, 487 GO terms, and 232 miRNAs. Disease features were characterized through disease-gene associations and disease-miRNA associations: disease-gene association data were obtained from the DisGeNET [23]; disease-miRNA association data were downloaded from the HMDD [24]. After deduplication and removal of all-zero entries, we obtained association information for 412 diseases, 10,146 genes, and 495 miRNAs.

To validate the cross-scenario application capability of our proposed method, we further collected drug-related association data for extended experiments. Drug-gene interaction data were downloaded from DGIdb [25]; drug-miRNA association data were obtained from SM2miR [26] and miR2Disease [27]; lncRNA-drug association data were collected from the LncDrug [28]. After data preprocessing, we obtained association information for 141 drugs, with drug features constructed through drug-gene and drug-miRNA associations, forming 2,068-dimensional feature vectors. These drug-related data will be used in subsequent cross-scenario application experiments to demonstrate the effectiveness of the model in lncRNA-drug and disease-drug association prediction tasks.

Let the lncRNA set be *L =* {*l*_1_,*l*_2_,…,*l*_*n*_}, the disease set be *D =* {*d*_1_,*d*_2_,…,*d*_*m*_}, and the drug set be 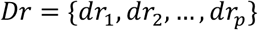. The heterogeneous feature matrix for lncRNAs is constructed by concatenating multiple types of biological association information:

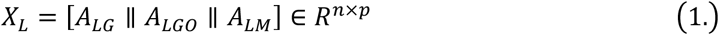

Where 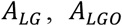 and 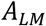 represent lncRNA-gene, lncRNA-GO, and lncRNA-miRNA association matrices, respectively, ∥ denotes matrix concatenation operation, resulting in 6,066-dimensional lncRNA features. Similarly, the disease feature matrix is constructed as:

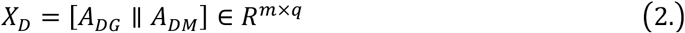

where *A*_*DG*_ and *A*_*DM*_ represent disease-gene and disease-miRNA association matrices, respectively, resulting in 10,641-dimensional disease features. The drug feature matrix is constructed as:

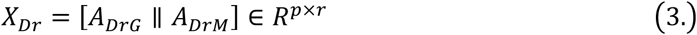

where *A*_*DRG*_ and *A*_*DRM*_ represent drug-gene and drug-miRNA association matrices, respectively, resulting in 2,068-dimensional drug features.

### 2.3 PCA-based Feature Dimensionality Reduction and Denoising

High-dimensional biomedical data commonly suffers from feature redundancy and noise issues, which not only increase computational complexity but may also lead to the curse of dimensionality and overfitting phenomena. Principal Component Analysis projects original high-dimensional data onto a low-dimensional subspace through linear transformation while preserving the main variational information of the data. For an input feature matrix *X* ∈ *R*^*N*×*D*^, where *N* is the number of samples and *D* is the feature dimension, we first center the data:

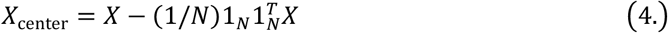

The covariance matrix is computed as:

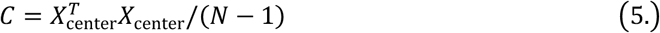

Principal components are obtained through eigenvalue decomposition:

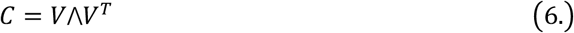

where *V =* [*v*_1_,*v*_2_,…,*v*_*D*_] is the eigenvector matrix, Λ *=* diag(λ_1_,λ_2_,…,λ_*D*_) is the eigenvalue diagonal matrix, and λ_1_ ≥ λ_2_ ≥ ⋯ ≥ λ_*D*_. The first *k* principal components are selected for dimensionality reduction

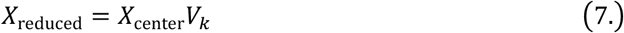

where *V*_*K*_ *=* [*v*_1_,*v*_2_,…,*v*_*k*_] contains the first *k* principal components. The selection of *k* is based on the cumulative contribution ratio criterion:

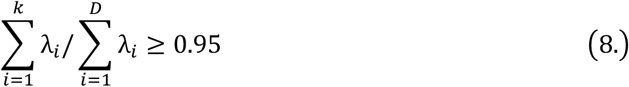

to retain 95% of the information. After PCA processing, we obtain the dimensionality-reduced feature matrices 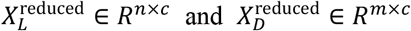, where *c* is the feature dimension after reduction.

### 2.4 Node Local Feature Encoding Based on Graph Attention Network

Graph Attention Networks address the limitation of traditional graph convolutional networks that treat all neighboring nodes with equal weights by introducing attention mechanisms. GAT can adaptively assign different importance weights to different neighboring nodes, thereby better capturing complex relationships between nodes. Based on the known lncRNA-disease association matrix *A* ∈ {0,1}^*n*×*m*^, we construct a heterogeneous bipartite graph *G =* (*V,E*), where the node set *V = V*_*L*_ ∪ *V*_*D*_, with *V*_*L*_ and *V*_*D*_ representing lncRNA and disease node sets, respectively; the edge set *E* contains all known lncRNA-disease associations. The PCA-processed features are used as initial node features:

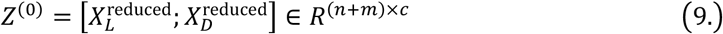

The adjacency matrix is constructed as:

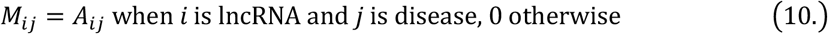

For any two connected nodes *i* and *j* in the graph, we first obtain high-order feature representations through linear transformation:

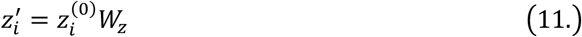

Where 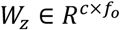 is a learnable weight matrix and *f*_*0*_ is the output feature dimension. The attention coefficients are computed as:

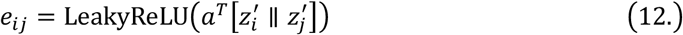

where 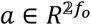 is the attention parameter vector. Attention weights are obtained through softmax normalization:

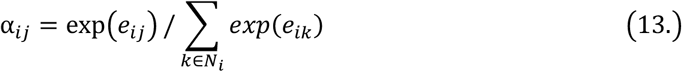

where 𝒩_*i*_ represents the neighbor set of node *i*. To stabilize the training process and enhance model expressiveness, a multi-head attention mechanism is adopted:

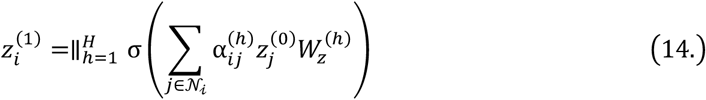

where *H* is the number of attention heads, σ is the ReLU activation function, and 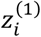 is the node representation after GAT encoding.

### 2.5 Multi-scale High-order Feature Fusion Based on MACM

Traditional graph neural networks primarily aggregate first-order neighbor information, suffering from limited receptive field problems and difficulty in effectively capturing long-range high-order structural relationships. In biological networks, associations between diseases and lncRNAs are often manifested through complex multi-hop paths, such as “lncRNA regulates genes → genes associate with diseases” or “lncRNA interacts with miRNAs → miRNAs regulate target genes → genes associate with diseases” and other multi-level regulatory relationships. The Multi-scale Adaptive Convolution Module (MACM) can comprehensively model multi-level topological structures from local to global by simultaneously considering neighborhood information of different hop counts, effectively addressing the over-smoothing problem in graph neural networks.

MACM takes the GAT output 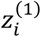 as input and computes node representations of different hop counts through a recursive approach. The 0-hop representation is initialized as the GAT output:

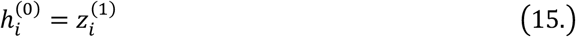

For the *h*-th hop (*h* ≥ 1), the node representation update formula is:

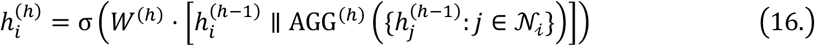

Where 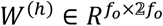 is the learnable weight matrix for the *h*-th hop, and ∥ denotes feature concatenation operation. The aggregation function employs mean aggregation:

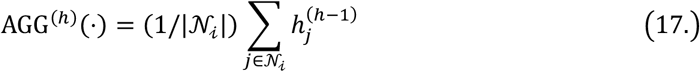

This design ensures that each hop’s representation contains both the node’s own information and neighborhood aggregated information, effectively preventing information loss. Information captured by different hop counts has different biological meanings: 1-hop neighborhoods reflect direct association relationships, representing known lncRNA-disease associations; 2-hop neighborhoods reveal indirect association patterns, such as potential connections established through commonly regulated genes or miRNAs; higher hop counts can discover more complex high-order regulatory network patterns.

To effectively integrate information from different scales, MACM employs an adaptive attention mechanism to compute importance weights for each scale. For node $i$, its adaptive weight is computed as:

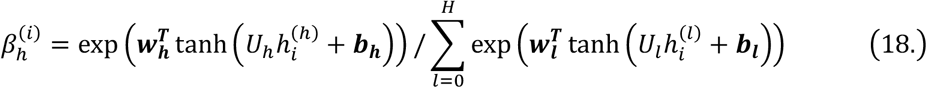

Where 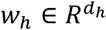 is the attention parameter vector, 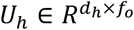 is the linear transformation matrix,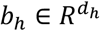 is the bias vector, and *d*_*h*_ is the attention hidden layer dimension. The final multi-scale node representation is obtained through weighted fusion:

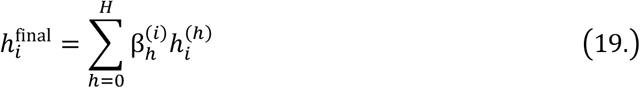

The advantages of this adaptive mechanism are threefold: first, different nodes can adaptively adjust the weights of different scale information according to their position and functional characteristics in the network, for example, central nodes may rely more on local information while peripheral nodes may need more global information; second, compared to simple averaging or max pooling, the attention mechanism can learn more refined scale importance patterns, improving the model’s expressiveness; finally, through the gating mechanism, the model can dynamically balance local accuracy and global coverage, avoiding the problem of long-range information diluting local features.

### 2.6 Association Prediction and Robustness Validation

For a given lncRNA-disease pair (*l*_*i*_,*d*_*j*_), we concatenate their corresponding multi-scale node representations:

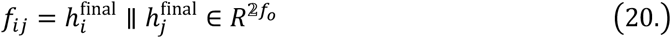

Non-linear transformation and association probability prediction are performed through a multi-layer perceptron:

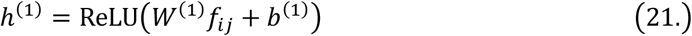

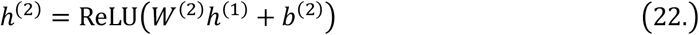

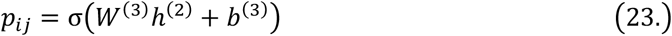

where σ is the sigmoid activation function, ensuring output probabilities are within the [0,1] range. The model is trained using a binary cross-entropy loss function:

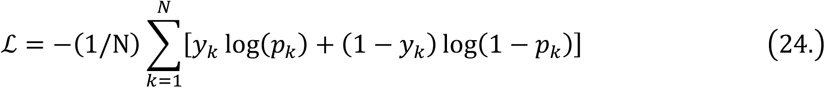

where *y*_*k*_ ∈ {0,1} is the true label and *p*_*k*_ is the predicted probability. To prevent overfitting, an L2 regularization term is added:

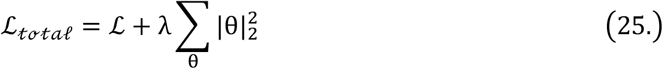

where λ is the regularization coefficient and θ represents all learnable parameters.

To validate the model’s robustness, we designed a perturbation strategy with random shuffling of feature columns. Specifically, we randomly permute the columns of the input feature matrix while maintaining the relative relationships between samples:

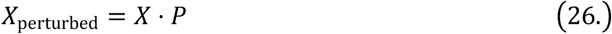

where *P* is a random permutation matrix. Multiple independent prediction experiments are conducted on each perturbed dataset, and the final prediction results are obtained through the following ranking aggregation strategy:

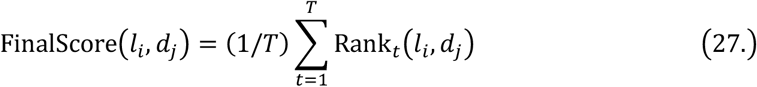

where *T* is the number of independent experiments and *R*a*nk*_*t*_(*l*_*i*_,*d*_*j*_) is the ranking of pair (*l*_*i*_,*d*_*j*_) in the *t*-th experiment. This strategy can effectively reduce the influence of random factors and improve the reliability of prediction results.

### 2.7 Disease-Drug Association Prediction Based on lncRNA Bridging

To further validate the application potential of our proposed method, we designed a prospective study that predicts disease-drug associations through lncRNAs as intermediate bridges. This method is based on “disease-lncRNA-drug” multi-hop association paths, first using the trained model to predict association probabilities between diseases and all lncRNAs, then inputting the drug feature matrix *X*_*Dr*_ into the same framework to predict lncRNA-drug associations. Specifically, for a given disease *d*_*i*_, we select the top-*K* ranked lncRNAs by predicted association probability as the candidate bridging node set 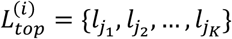. Subsequently, we predict the association probabilities between these candidate lncRNAs and all drugs. The final association score between disease *d*_*i*_ and drug 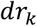 is calculated through the maximum association path via bridging lncRNAs:

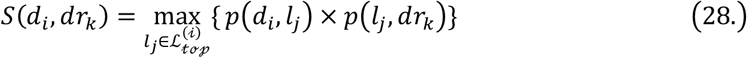

where *p*(*d*_*i*_,*l*_*j*_) and 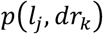 represent the predicted association probabilities for disease-lncRNA and lncRNA-drug, respectively. This lncRNA bridging strategy can not only discover potential disease-drug associations but also provide new research directions for drug repositioning and precision medicine. By utilizing drug association information from DGIdb, SM2miR, miR2Disease, and LncDrug databases, this method demonstrates the broad applicability of the hybrid neural network framework in cross-biomolecular type association prediction.

## 3. Experiments and results

### 3.1 Evaluation criteria

To comprehensively assess the predictive performance of our model, we employed a five-fold cross-validation strategy. Specifically, the set of all known lncRNA-disease association samples was randomly partitioned into five equal subsets; in each iteration, one subset served as the test set while the remaining four formed the training set. This procedure was repeated five times and the resulting performance metrics were averaged. For each fold, we measured the model’s discriminative ability using the area under the receiver operating characteristic curve (AUC). Let {*s*_*p*_}and {*s*_*n*_} denote the predicted scores for the positive (associated) and negative (non-associated) samples in a given fold, and let 𝒫 and 𝒩 be the corresponding index sets. The AUC is defined as

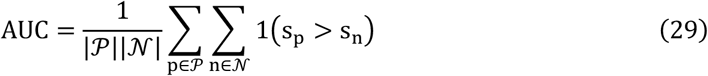

where1(·) is the indicator function, equal to 1 if *s*_*p*_ > *s*_*n*_ and 0 otherwise.

Furthermore, we plotted Precision-Recall (PR) curves to evaluate the model’s performance under class imbalance scenarios, calculating the corresponding area under the PR curve (AUPR):

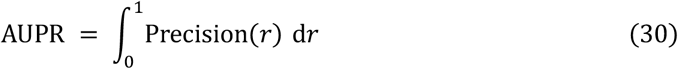

Finally, we obtained the average metrics across the five folds, defined as:

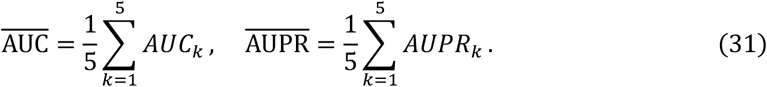

### 3.2 Five-Fold Cross-Validation

In the five-fold cross-validation experiments, we compared the proposed MACGA model with four representative methods: DMFLDA [29], GCLMTP [30], LRGNN [31], and GANLDA [32]. ROC curves (Figure 2) and PR curves (Figure 3) were plotted to evaluate the predictive performance of each model. As shown in Figure 2, MACGA achieved the highest AUC value (0.9463), significantly outperforming DMFLDA (0.8854), GCLMTP (0.9366), LRGNN (0.9118), and GANLDA (0.8881). Furthermore, in the region with a low false positive rate (FPR<0.1), the ROC curve of MACGA consistently stayed above those of the other four methods.

**Fig 2.**
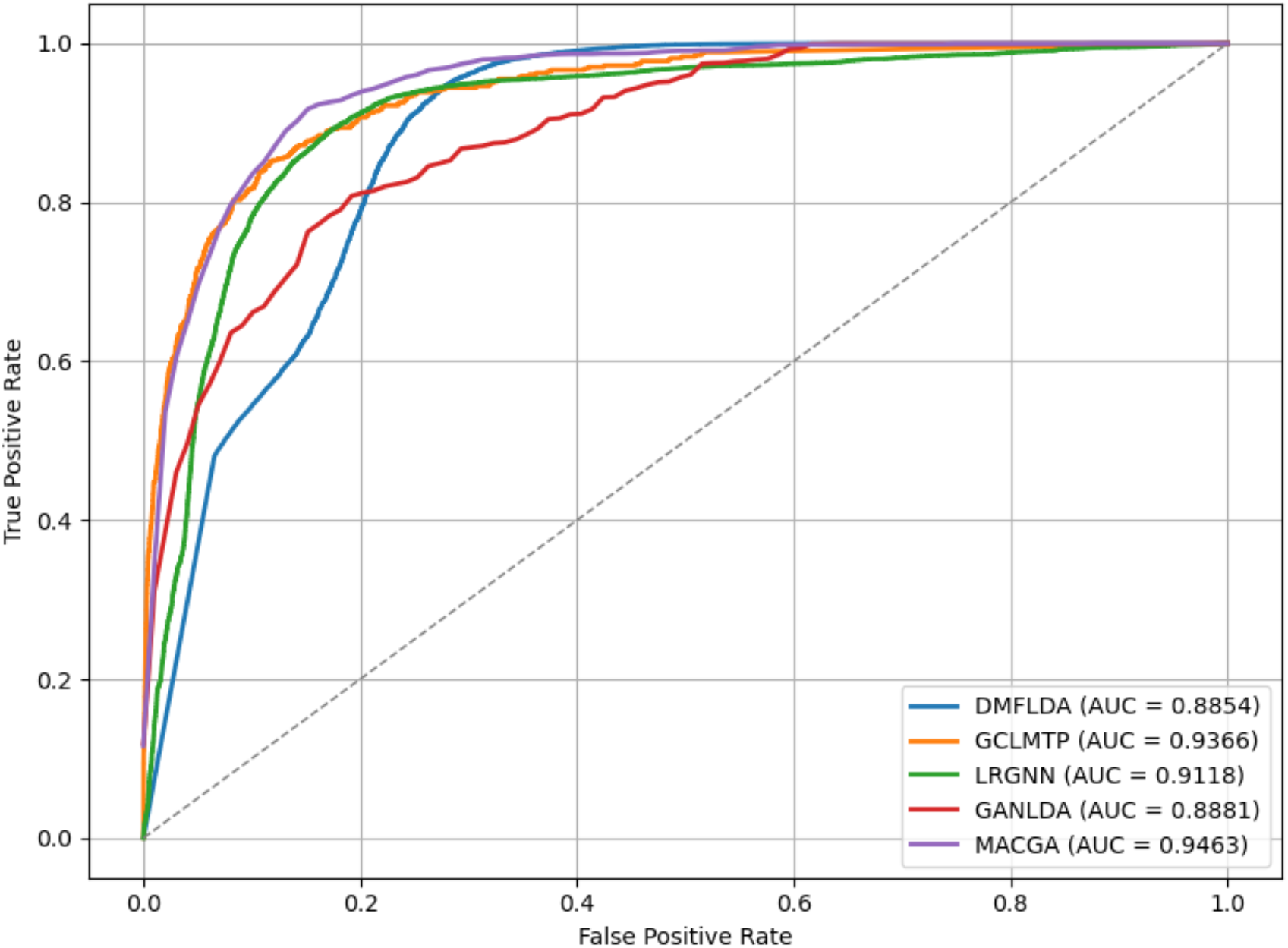
The performance comparison of MACGA, GANLDA, LRGNN, GCLMTP and DMFLDA in term of AUC value based on 5-fold cross validation.

**Fig 3.**
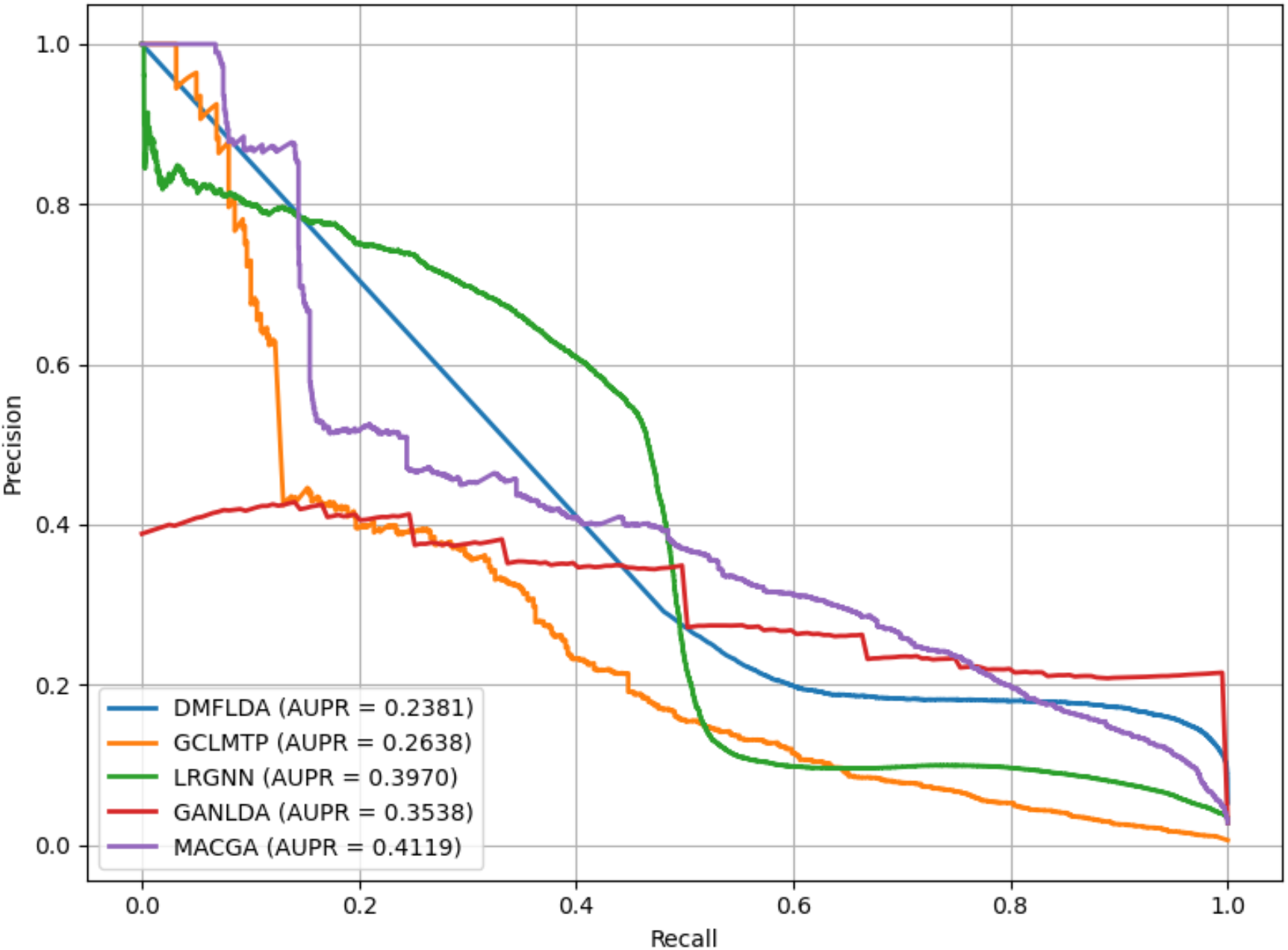
The performance comparison of MACGA,GANLDA,LRGNN,GCLMTP and DMFLDA in term of AUPR value based on 5-fold cross validation.

Considering the imbalance of positive and negative samples in the dataset, we further plotted Precision-Recall (PR) curves to evaluate the performance of each method. As illustrated in Figure 3, MACGA similarly attained the highest AUPR value (0.4119), markedly exceeding DMFLDA (0.2381), GCLMTP (0.2638), LRGNN (0.3970), and GANLDA (0.3538). These results indicate that the MACGA model effectively balances precision and recall under class-imbalanced conditions.

### 3.3 Parameter setting

To ensure optimal model performance, we employed a staged grid search strategy to systematically optimize key hyperparameters across all modules on the dataset. For the PCA dimensionality reduction module, the optimal feature dimension was determined to be 128 through comparative validation of different dimensional candidates. In the Graph Attention Network (GAT) module, testing of various dropout rates revealed that 0.4 most effectively balances overfitting prevention and information retention. For the Multi-scale Adaptive Convolutional Module (MACM), systematic optimization established the optimal configuration with an input dimension of 64 and sigma parameter of 0.9. In the Multi-Layer Perceptron (MLP) prediction layer, comparative testing resulted in the selection of 64 as the number of hidden units in the final layer. Regarding training hyperparameters, grid search within the learning rate range {0.05-0.00005} and weight decay values {1×10^−5^, 5×10^−5^, 1×10^−4^, 5×10^−4^, 5×10^−3^} identified the optimal combination as a learning rate of 0.006 and weight decay of 1×10−^4^. All hyperparameter configurations were validated through multiple rounds of experiments, significantly enhancing model stability and prediction accuracy.

### 3.4 Case studies

To further evaluate the predictive performance and generalization capability of the MACGA model for disease-related lncRNAs, we conducted case studies on two representative diseases, prostate cancer and colon cancer. Specifically, the MACGA model was first trained using all known lncRNA-disease association data. Subsequently, predictions of unknown disease-associated lncRNAs were generated as candidate associations, which were ranked according to predicted probabilities. These predicted associations were then individually verified through literature searches in the PubMed database. Additionally, to further assess the generalization and practical applicability of the MACGA model, we extended our predictions from disease-lncRNA associations to disease-drug associations, evaluating the feasibility and accuracy of the MACGA approach for drug repositioning.

As shown in Table 1, for prostate cancer, all of the top 20 lncRNAs predicted by the MACGA model (20/20) have been experimentally validated in previous studies. When the list is expanded to the top 30 candidates, 27 lncRNAs are explicitly supported by literature reports in PubMed, with only three lncRNAs (BANCR, LSINCT5, and ESRG) lacking direct experimental evidence. Notably, the highest-ranked lncRNAs, including H19 (PMID:32356434), CDKN2B-AS1 (PMID:39632396), HOTAIR (PMID:33024272), MEG3 (PMID:38047026), and MALAT1 (PMID:33428963), have been consistently reported to play crucial regulatory roles in the initiation and progression of prostate cancer. These results underscore the precision and reliability of the MACGA model in identifying high-confidence disease-associated lncRNAs.

**Table 1.** The top 30 predicted results of prostate cancer-related lncRNAs based on MACGA.

| Rank | LncRNA name | Evidence | Rank | LncRNA name | Evidence |
| --- | --- | --- | --- | --- | --- |
| 1 | H19 | PMID:32356434 | 16 | CCAT2 | PMID:32831916 |
| 2 | CDKN2B-AS1 | PMID:39632396 | 17 | HULC | PMID:35178357 |
| 3 | HOTAIR | PMID:33024272 | 18 | SPRY4-IT1 | PMID:36909367 |
| 4 | MEG3 | PMID:38047026 | 19 | TP53COR1 | PMID:25999983 |
| 5 | MALAT1 | PMID:33428963 | 20 | AFAP1-AS1 | PMID:31669642 |
| 6 | PVT1 | PMID:30914434 | 21 | BANCR | unknown |
| 7 | GAS5 | PMID:37837861 | 22 | MIR155HG | PMID:28188157 |
| 8 | UCA1 | PMID:32805084 | 23 | LINC00675 | PMID:32801300 |
| 9 | NEAT1 | PMID:39379919 | 24 | LSINCT5 | PMID:unknown |
| 10 | MIR17HG | PMID:26891588 | 25 | BCYRN1 | PMID:32705287 |
| 11 | HOTTIP | PMID:29884766 | 26 | HCP5 | PMID:31746434 |
| 12 | KCNQ1OT1 | PMID:33688211 | 27 | IGF2-AS | PMID:25670076 |
| 13 | TUG1 | PMID:31210308 | 28 | WT1-AS | PMID:34504167 |
| 14 | CCAT1 | PMID:34319909 | 29 | DLEU2 | PMID:35075115 |
| 15 | XIST | PMID:36038831 | 30 | ESRG | unknown |

As detailed in Table 2, for colon cancer, 18 of the top 20 predicted lncRNAs have literature support, with only BANCR and LINC00675 lacking relevant reports. When extending the predictions to the top 30, a total of 27 lncRNAs have been validated, and only three lncRNAs (BANCR, LINC00675, and KIRREL3-AS3) remain unreported. Among the highest-ranked lncRNAs, HOTAIR (rank 1, PMID:30541551) has been reported to significantly promote colon cancer cell proliferation and invasion. Similarly, MEG3 (rank 2, PMID:39108882) and MALAT1 (rank 3, PMID:31733641) have been documented as regulators of critical oncogenic pathways in colorectal cancer. These findings further confirm the robustness and accuracy of the MACGA model in predicting disease-associated lncRNAs across diverse disease contexts.

**Table 2.** The top 30 predicted results of colon cancer-related lncRNAs based on MACGA.

| Rank | LncRNA name | Evidence | Rank | LncRNA name | Evidence |
| --- | --- | --- | --- | --- | --- |
| 1 | HOTAIR | PMID:30541551 | 16 | SPRY4-IT1 | PMID:28651500 |
| 2 | MEG3 | PMID:39108882 | 17 | HULC | PMID:30551459 |
| 3 | MALAT1 | PMID:31733641 | 18 | GHET1 | PMID:27131316 |
| 4 | H19 | PMID:28358427 | 19 | AFAP1-AS1 | PMID:30588252 |
| 5 | UCA1 | PMID:30652355 | 20 | LINC00675 | unknown |
| 6 | GAS5 | PMID:27863421 | 21 | SNHG12 | PMID:34650693 |
| 7 | CDKN2B-AS1 | PMID:32767927 | 22 | CASC2 | PMID:32655801 |
| 8 | HOTTIP | PMID:29274585 | 23 | XIST | PMID:29679755 |
| 9 | CCAT2 | PMID:28964256 | 24 | TUG1 | PMID:27634385 |
| 10 | NEAT1 | PMID:32077635 | 25 | KIRREL3-AS3 | unknown |
| 11 | MIR17HG | PMID:35116852 | 26 | HNF1A-AS1 | PMID:35184384 |
| 12 | PVT1 | PMID:35763386 | 27 | ZEB1-AS1 | PMID:35794981 |
| 13 | BANCR | unknown | 28 | LINC01133 | PMID:27443606 |
| 14 | TUSC7 | PMID:31002365 | 29 | TINCR | PMID:37051356 |
| 15 | CCAT1 | PMID:25185650 | 30 | BCYRN1 | PMID:31773686 |

Moreover, we applied the MACGA model in combination with the proposed lncRNA-bridging strategy to predict potential therapeutic drugs for prostate cancer, aiming to explore its applicability for drug repositioning. As shown in Table 3, among the top 10 drugs predicted for prostate cancer, seven have already been validated by clinical or preclinical studies according to PubMed references. For example, the top-ranked drug, 17-aag (PMID:24130883), has been experimentally demonstrated to exhibit significant anti-cancer activity against prostate cancer cells. Similarly, the second-ranked drug, sorafenib (PMID:31037974), has demonstrated promising therapeutic effects in clinical investigations. Collectively, these results indicate that the proposed MACGA approach can not only accurately predict disease-associated lncRNAs but also holds substantial potential for identifying novel therapeutic agents and facilitating drug repositioning research.

**Table 3.**
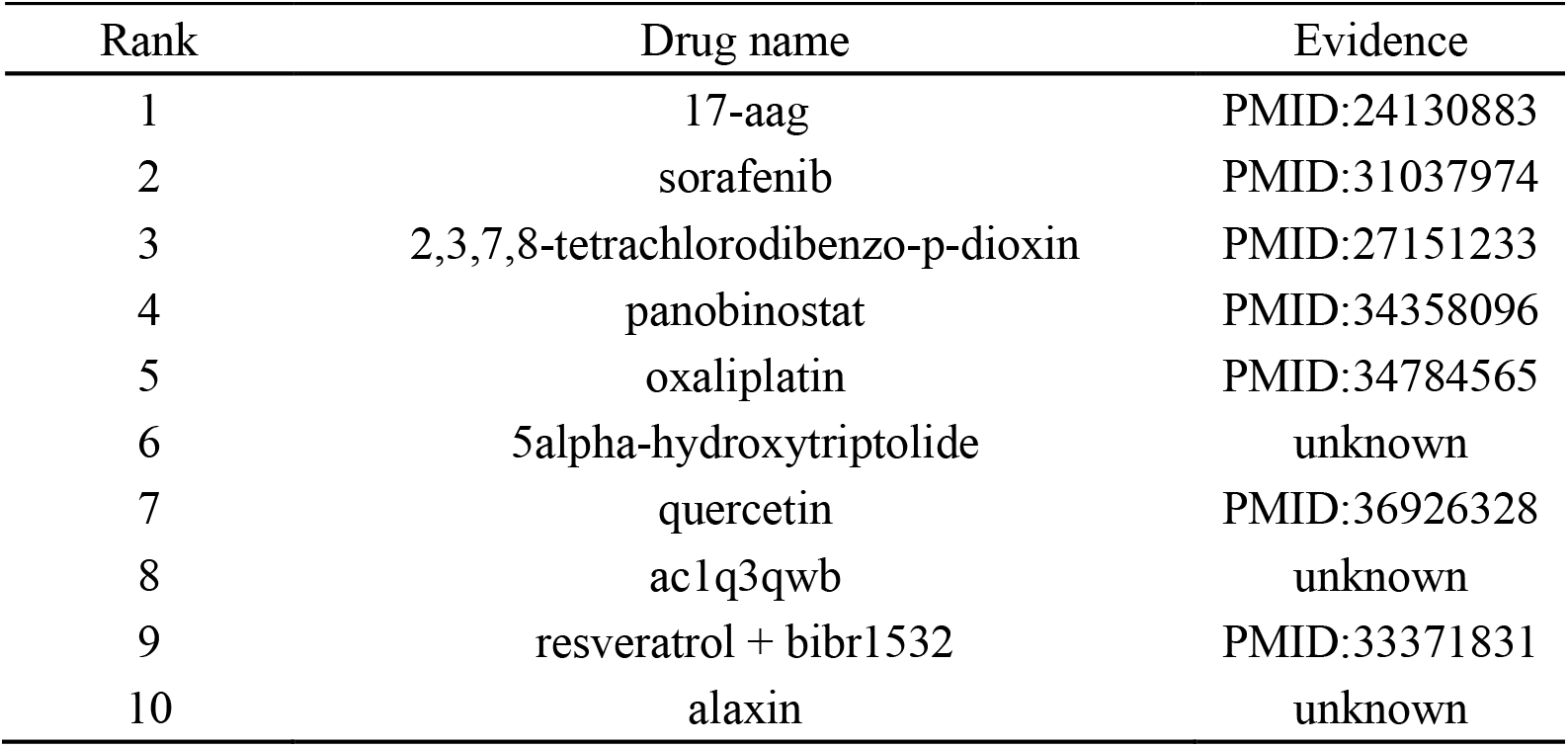
The top 10 predicted results of prostate cancer-related Drugs based on MACGA.

## 4. Conclusion

In this study, we propose a hybrid neural network framework termed MACGA, which integrates Graph Attention Networks (GAT) and a Multi-scale Adaptive Convolution Module (MACM) to address the oversmoothing problem frequently encountered in deep graph neural network (GNN) models. MACGA achieves high-accuracy predictions of lncRNA-disease associations through an end-to-end pipeline involving PCA-based feature dimensionality reduction, local feature extraction via GAT, and multi-scale feature fusion by MACM. Experimental results indicate that MACGA significantly outperforms four representative comparison methods. Moreover, case studies conducted on prostate cancer and colon cancer validated the reliability and precision of MACGA’s predictions. We also successfully extended the MACGA framework to predict disease-drug associations, demonstrating its promising application in drug repositioning and precision medicine.

Future studies could further improve the generalizability of MACGA by incorporating additional heterogeneous biomedical data sources and investigate more deeply into its potential applications in drug repositioning research.

